# Dental ontogeny in the most primitive bony fish *Lophosteus* reveals the developmental relationship between teeth and dermal odontodes

**DOI:** 10.1101/2020.07.14.202234

**Authors:** Donglei Chen, Henning Blom, Sophie Sanchez, Paul Tafforeau, Tiiu Märss, Per E. Ahlberg

## Abstract

Ontogenetic data obtained by synchrotron microtomography of a marginal jawbone of *Lophosteus superbus* (Late Silurian, 422 Million years old), the phylogenetically basalmost stem osteichthyan, reveal developmental relationships between teeth and ornament that are not obvious from the adult morphology. The earliest odontodes are two longitudinal founder ridges formed at the ossification center. Subsequent odontodes that are added lingually to the ridges turn into conical teeth and undergo cyclic replacement, while those added labially achieve a stellate appearance. The stellate odontodes deposited directly on the bony plate are aligned with the alternate files of the teeth. Successive odontodes overgrowing the labial tooth rows become tooth-like and the replacement teeth near to them are ornament-like. We propose that teeth and ornament are modifications of a single odontode system regulated and differentiated by the oral and dermal signals; signal cross-communication between the two domains can occur around the oral-dermal boundary.

## Introduction

A tooth is a particular type of odontode, the characteristic integrated unit formed by dental tissues (Huysseune and Sire, 1998). Although teeth are the only odontodes to persist in tetrapods, various forms of dermal odontodes covering the outer surface of the body are present in certain jawed fishes as well as in the extinct jawless vertebrates known as “ostracoderms” (Janvier, 1996). Probably the most familiar examples of dermal odontodes are the placoid scales of sharks, which appear so similar to the teeth that they are sometimes referred to as “skin teeth”. On one hand, teeth and dermal odontodes are considered as homologous structures from the same developmental module as evidenced by a common gene regulatory network (Debiais-Thibaud et al., 2011). On the other hand, they are regarded as separated systems because dermal odontodes lack the regular temporo-spatial pattern of teeth, based on the observations from modern sharks (Reif, 1982; Fraser and Smith, 2011). However, the growth patterns of dermal odontodes are generally poorly understood.

The evolutionary developmental relationship between teeth and dermal odontodes is pivotal for understanding the origin of teeth. The classic “outside-in” hypothesis is currently in favor after decades of debate (Donoghue and Rücklin, 2014), but the developmental continuum between teeth and dermal odontodes, which is one of its central premises, still lacks unequivocal evidence. Extant gnathostomes with dermal odontodes (sharks, rays and some bony fishes such as *Polypterus)* always display a sharp demarcation between teeth and ornament. Even though they can provide data of all ontogenetic stages, they are not informative about the evolution of the developmental relationship between teeth and dermal odontodes. For that we must turn to the fossil record of the earliest jawed vertebrates, in particular to the jawed stem gnathostomes, which form the common ancestral stock of Chondrichthyes + Osteichthyes, and the stem osteichthyans, which form the common ancestral stock of Actinopterygii + Sarcopterygii (Figure 1). This paper presents the marginal dentition of the Late Silurian (422 million years old; https://stratigraphy.org/timescale/) stem osteichthyan *Lophosteus superbus,* based on investigation by propagation phasecontrast synchrotron microtomography (PPC-SRμCT), which allows the dentition to be digitally dissected in 3D with sub-micrometer resolution.

**Figure 1.**
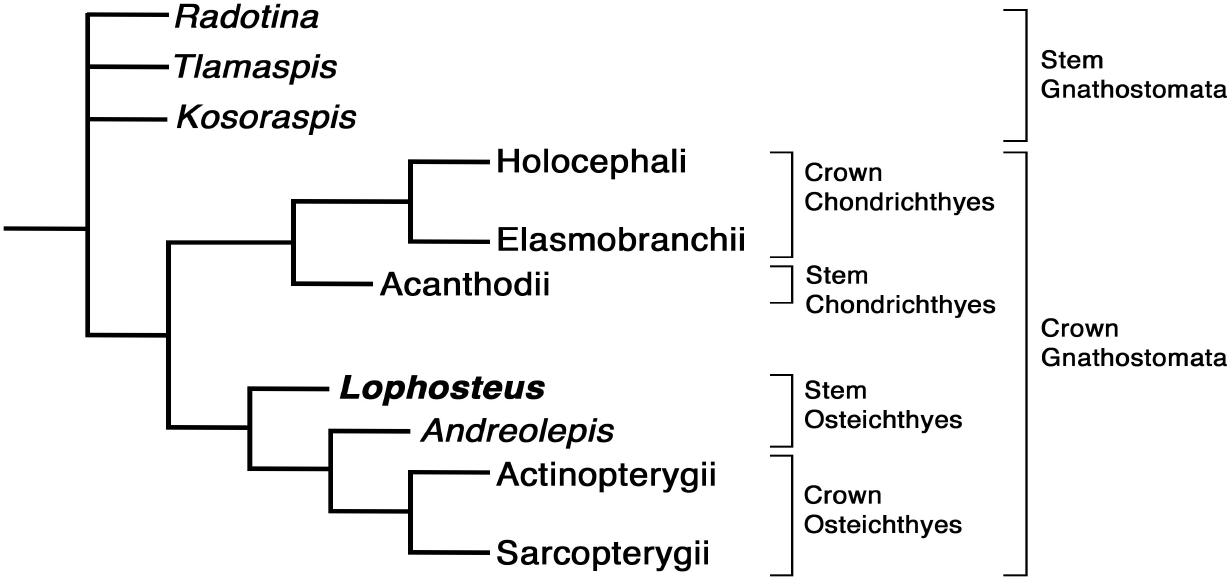
Phylogenetic position of *Lophosteus* and some of the other fossil taxa discussed in this paper. Tree topology, from Qu et al. (2015) and Vaškaninová et al. (2020). Formal hierarchical categories indicated on right.

The same technique has revealed the most primitive gnathostome dentitions (Vaškaninová et al., 2020) and the earliest osteichthyan-style tooth replacement (Chen et al., 2016). The phylogenetically most basal vertebrates known to possess teeth are the Early Devonian armoured fish known as ‘acanthothoracids’, including *Radotina, Kosorapis* and *Tlamaspis* (Figure 1). All their dentitions suggest that being added lingually but not shed and solely carried by marginal dermal bones are the ancestral conditions of teeth (Vaškaninová et al., 2020). Chondrichthyans and osteichthyans both inherit the lingual tooth addition, but evolve tooth shedding independently; while marginal jawbone ornamented by dermal odontodes is only kept by osteichthyans. The jawbones of *Kosoraspis* and *Tlamaspis* consist of multiple short pieces (Vaškaninová et al., 2020, fig. 3–4), a condition strikingly retained in *Lophosteus* (Figure 2), but unknown in other taxa. *Kosoraspis* show no sharp demarcation between teeth and ornament. *Kosoraspis*-like tooth-ornament gradation is combined with cyclic tooth shedding in *Lophosteus* (Botella et al., 2007; Chen et al., 2017), as well as in the oldest known stem osteichthyan *Andreolepis* (424 million years old; Chen et al., 2016). *Radotina* displays stellate teeth that are extremely similar to the stellate dermal odontodes, which is the characteristic ornament of ‘acanthothoracids’. Interestingly, the ornament of *Lophosteus,* but not *Andreolepis,* share a distinctive stellate morphology with ‘acanthothoracids’ (Burrow, 1995). *Lophosteus* also resembles a stem gnathostome in completely lacking enamel, whereas *Andreolepis* has enamel on its scales (Qu et al., 2015). This character distribution suggests that *Lophosteus* is the least crownward of known stem osteichthyans (Figure 1), making it uniquely informative about the evolution of the osteichthyan dentitions in conjunction with the ‘acanthothoracid’ dentitions.

**Figure 2.**
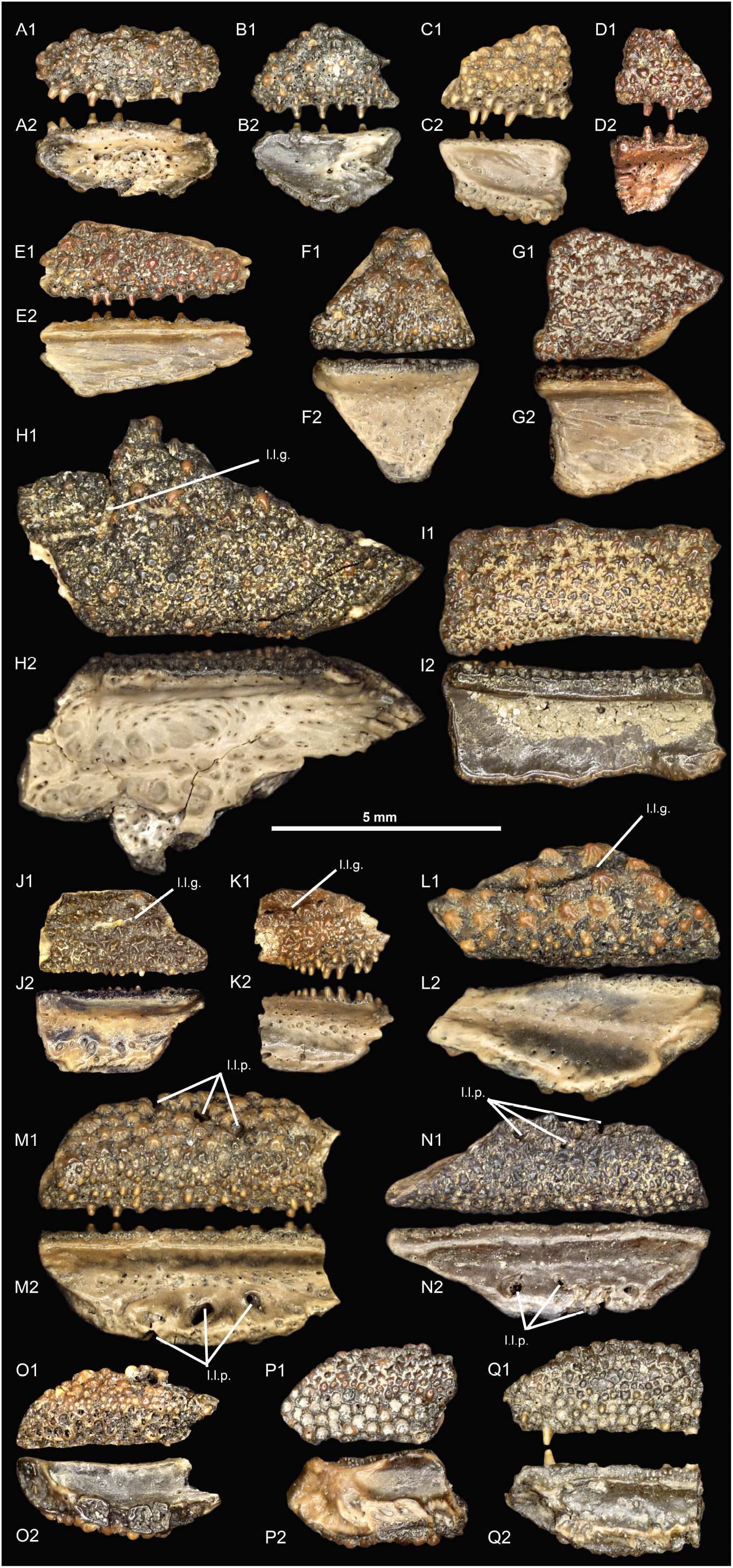
Morphological variation of the marginal jawbones in *Lophosteus.* Scale bar, 5 mm. A, GIT 760-13, B, GIT 760-14, C, GIT 760-15, D, GIT 760-16, E, GIT 760-17, F, GIT 760-18, G, GIT 760-19, H, GIT 760-20, I, GIT 760-21, J, GIT 760-12, K, GIT 760-22, L, GIT 760-23, M, GIT 760-24, N, GIT 760-25, O, GIT 760-26, P, GIT 760-27, Q, GIT 760-28. 1, external view; 2, visceral view. l.l.g., lateral line groove.

## Results

A single marginal jawbone of *Lophosteus,* GIT 760-12 (Figure 2J) from the Late Silurian (Pridoli) of Ohessaare Cliff, Saaremaa, Estonia, was scanned with an isotropic voxel (3D pixel) size of 0.696μm. This specimen is probably from the lower jaw as it carries a lateral line groove that parallels the jaw margin. It can be approximately divided into oral and facial laminae, which form a transverse angle of about 50° (Figure 3 and 4A). The ossification center is likely located at the boundary of the two laminae, the thickest point of the bony plate (Figure 3). The feeder blood vessels for the odontode layer, which penetrate the bony plate with an increasing diameter and obliquity from lingual to labial, all radiate from here (Figure 3B, sky blue; Figure 5B and 6A, arrows).

**Figure 3:**
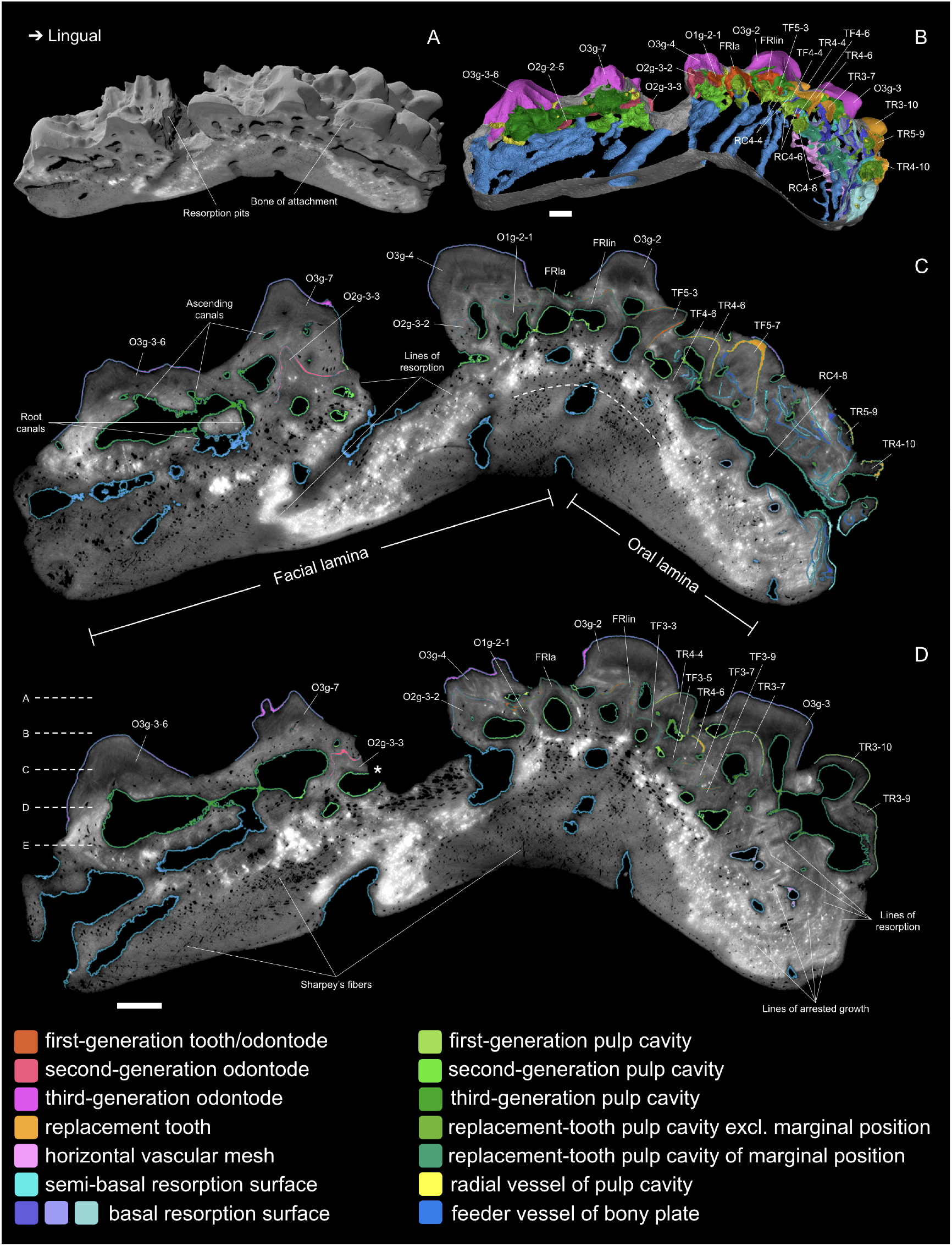
Virtual transverse sections between File 3 and 5. A, perspective block view from an antero-external angle. B, antero-visceral view of a slab of the 3D histological model through the midline of File 4. C-D, 2D sections. Resorption surfaces are not highlighted in D to show the preservation and scan quality. Dashed curve, the putative range of the jawbone when first mineralized, corresponding to the radiation center of the feeder vessels in B. Dashed lines, sectioning planes of Slideshow 1A-E. Asterisk, pulp cavity exposed by the extensive resorption during the development of the lateral line canal. Abbreviations and numbering rules see “Materials and Methods”. Scale bar, 0.1 mm.

**Figure 4.**
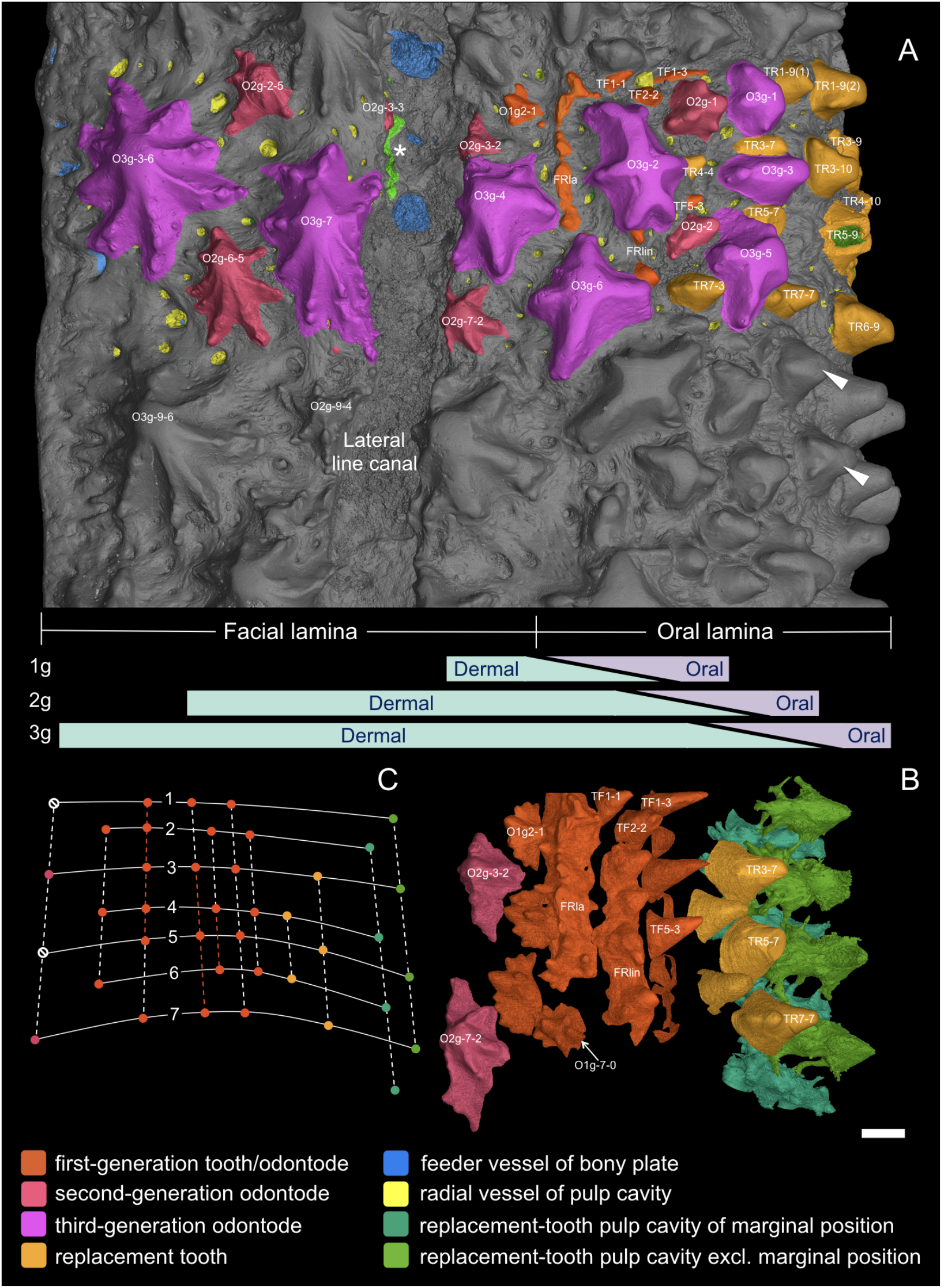
3D external view of scanned area of the marginal jawbone, the same view as the sections in Slideshow 1. Note that the jawbone is convex, see Figure 2. A, external morphology with modelled structures highlighted in colours, compared with the indistinguishability of the uncoloured area. Note, bone of attachment is not coloured, but the series of concentric arcs labial to teeth, e.g. TR5-9 and TR6-9, are edges of resorption surfaces that segregate the successive bone of attachment (see Figure 2B-C, TR5-9). Asterisk, exposed pulp cavity. Arrow head, more ornament-like teeth in the uncoloured area. Bars in lavender and mint, the putative gradient of the oral and dermal signal spheres, respectively, and the shift of oral-dermal boundary (overlap of the two signal spheres) along with the deposition of the first-generation (1g), second-generation (2g) and third-generation (3g) ornament odontodes. B, aligned with A, overgrowing odontodes and bone matrix are made invisible to show a consistent alternate pattern between replacement teeth, first-generation teeth and ornament odontodes lingual to the lateral line canal. The replacement teeth at the inserted positions are not shown. Because the lingual rows of first-generation teeth are obscured by the labial rows of replacement teeth in this view, these rows are not shown (for all tooth rows, see Figure 5A-B). For the best visibility, the most lingual rows of replacement teeth are represented by their pulp cavities. Except O1g-7-0, odontodes labelled with numbers are partially exposed and visible in A. C, diagram of the alternate organization based on B. Solid lines, transverse files; numbers of files mark the putative level of the ossification center. Dashed lines, longitudinal rows; colored parts of the dashed lines indicate the range of the founder ridges. Dots, positions of the structures in the particular colors. Null signs, suppressed positions. Scale bar, 0.1 mm.

The odontode component of the jaw bone consists of an extremely complex multilayered assemblage of ornament odontodes, teeth (complete or partly resorbed), pulp cavities, and resorption surfaces. The PPC-SRμCT data allow us to dissect this assemblage completely and reconstruct its entire ontogenetic history from the initiation of the first odontodes through to the death of the fish. The results and primary interpretations are presented here, beginning with the initiation of the initial odontodes and then going on to describe the teeth and dermal odontodes separately, followed by a consideration of the inferred developmental interaction between teeth and odontodes.

### The earliest odontodes

Two rows of first-generation odontodes are fused into longitudinal ridges with confluent pulp cavities at the level of the probable ossification center (Figure 4B and 5B, FRla, FRlin). These appear to be the ontogenetically earliest odontodes of the entire system (Figure 7A); we designate them as “founder ridges”. There are subtle differences between the two. On the lingual founder ridge, the main cusps are tall, conical, lingually recurved and widely spaced; on the labial founder ridge they are blade-like, labially inclined and united. In some places they are linked by small cusps. Side cusps are more numerous on the labial founder ridge. The labial flanks of both ridges carry more side cusps than the lingual flanks (Figure 6F). The lingual ridge ends posteriorly followed by unicuspid isolated odontodes (Figure 4B, TF1-1 and TF2-2. Odontode numbering see Materials and Methods), while the labial ridge ends anteriorly followed by a multicuspid isolated odontode (Figure 4B, O1g-7-0). The main cusps of the two founder ridges and isolated odontodes are aligned in rows (Figure 4B; Slideshow 1). There is no overlap between the two ridges; instead, their bases join like a continuous sheet (Figure 3B), implying they formed simultaneously.

**Figure 5.**
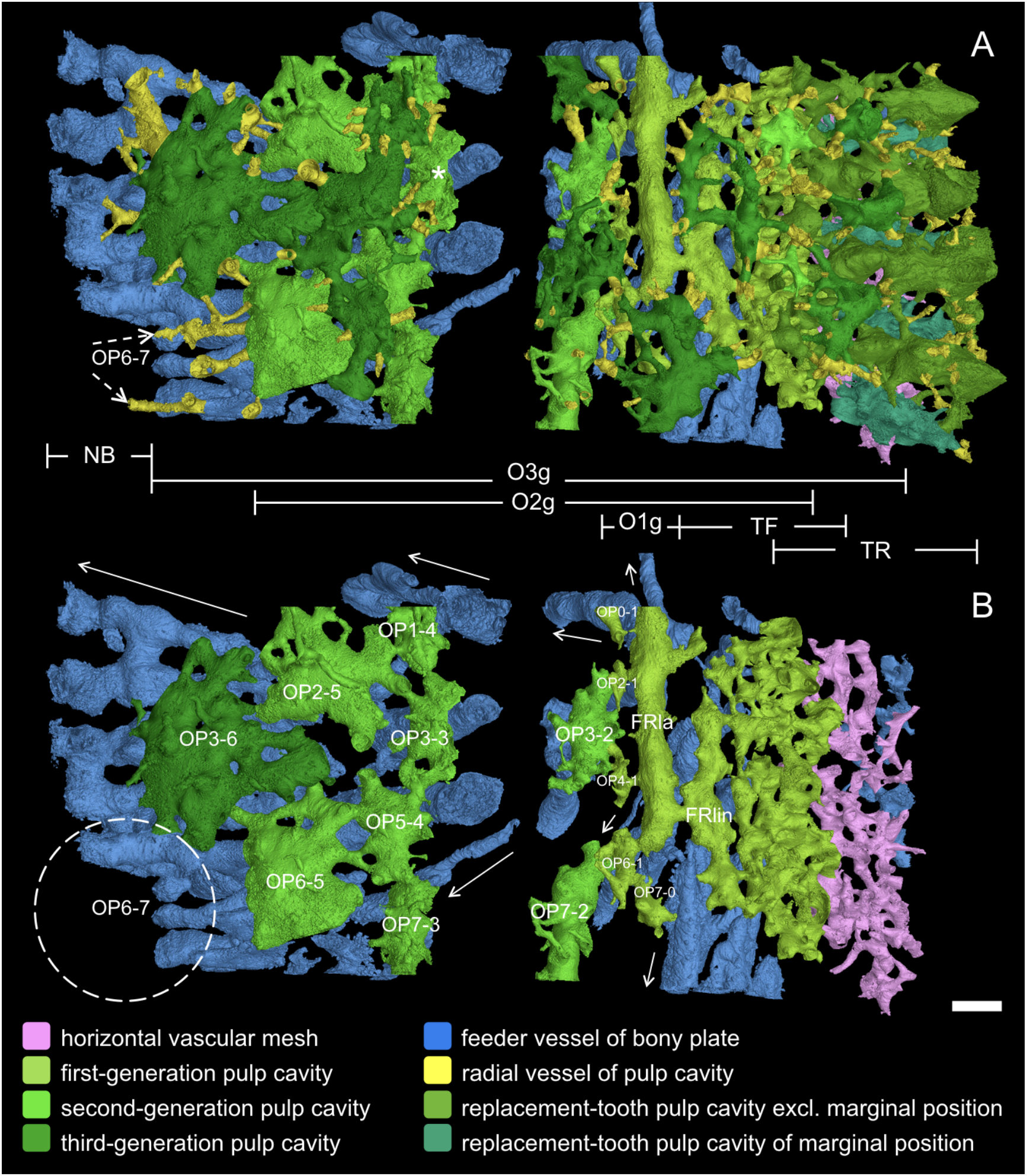
3D external views (same as Figure 3) of internal vascular system. A, all canals in the modelled area. Asterisk, exposed pulp cavity in Figure 4A and 7D. Dashed arrows, canals in yellow presumably growing from the developing odontode at OP6-7, see B. B, radial canals and pulp cavities of the overgrowing odontodes and replacement teeth are digitally removed to expose the pulp cavities that are laid down directly on the basal bone. The labelled numbers point out that they are organized in alternate rows, with some positions suppressed. Numbering of odontode positions (OP) is continued in Figure 5A,B. TR and TF, ranges of tooth replacement columns and first-generation teeth, indicating the putative drift of the oral epithelium. O1g, O2g and O3g, ranges of the first, second and third generations of ornament odontodes, indicating the expansion of the dermal epithelium. TF and 1g illustrate the range of the first mineralized part of bone, see Figure 6B. NB, new bone margin. Dashed oval, the next odontodes predicted to be added at the new labial margin. Arrows, the directions of the feeder vessels as they penetrate the bony plate from the ossification center, through the pulp cavities, to the openings on the internal surface and the lingual edge, see Figure 6A-B. Scale bar, 0.1 mm.

**Figure 6.**
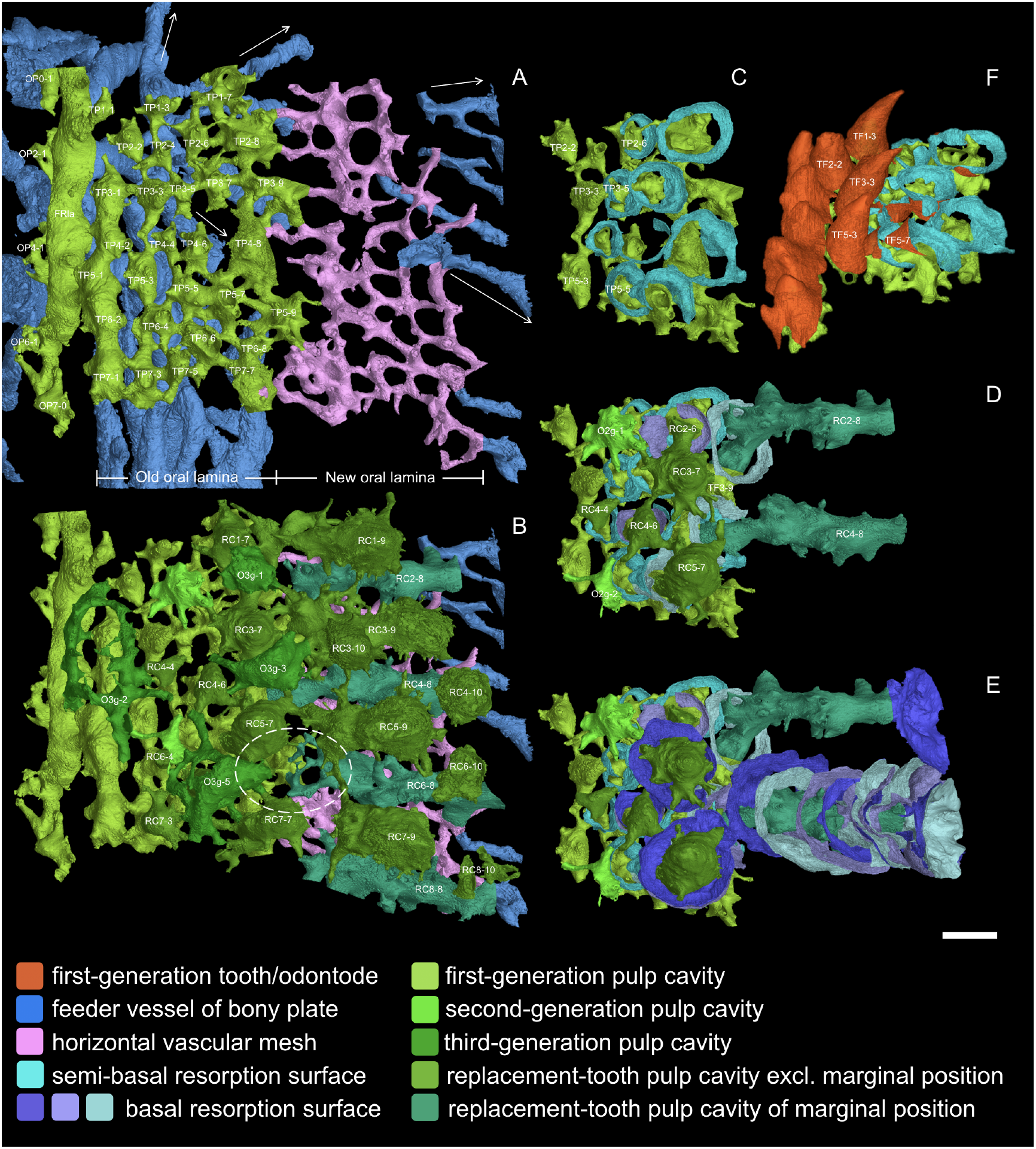
Occlusal view of the oral lamina. A-B, numbering of tooth positions (TP). A, pulp cavities of the first-generation teeth and odontodes. Arrows, the directions of the feeder vessels radiating from the ossification center, see Figure 5B. The feeder vessels in blue run longitudinally only beneath the first-generation teeth, but never do it labially, and these longitudinal feeder vessels may have penetrated the old oral lamina at the early stage. The horizontal vascular mesh in pink, which represents the new oral lamina beyond the first-generation teeth, supports the lingual replacement columns in the interval that lacks feeder vessels. B, pulp cavities of replacement teeth and overgrowing odontodes are added. For the feeder vessels, only those newly incorporated at the jaw margin are shown. Note that RC2-6 is covered by O3g-1, and RC6-6 is covered by O3g-5. Dashed oval, an example of the discontinuity of the pulp cavities between a first-generation tooth and its successive replacement teeth, which suggests the replacement of the first generation tooth has been delayed, but the drift of the belayed replacement teeth still follows the same file. C-E, comparison between the tooth replacement of File 2-3 and File 4-5. The successive resorption surfaces of RC2-4, which are similar to those of RC4-8, are not shown, except the last and the first basal resorption surface and the semi-basal resorption surface. Note that the resorption surfaces of RC4-8 gradually change in orientation. F, antero-occlusal view from lingual founder ridge to non-shedding and semi-basally resorbed first-generation teeth. Scale bar, 0.1 mm.

Following the establishment of the founder ridges, more isolated odontodes were added sequentially in both lingual and labial directions, overlapping the lingual and labial edges of the founder ridges, respectively (Figure 3B, O1g-2-1 and TF5-3). These new odontodes are unicuspid, conical teeth on the lingual side of the lingual ridge (Figure 6F), but are multicuspid and quickly take on the form of stellate tubercles with crenulated ridges on the labial side of the labial ridge (Figure 4B). Simply put, as the odontode skeleton spreads away from the two founder ridges, it turns into teeth lingually and into ornament labially.

### The Teeth

In the rows that follow lingually from the lingual founder ridge, all the main cusps become isolated and sharp. Although variable in size, they are arranged in semiregular alternate files, and the lingual founder ridge can be regarded as a union of two alternate rows (Figure 4B-C; Figure 6A, TP3-1, TP5-1, TP7-1 and TP4-2, TP6-2). These unicuspid odontodes are considered as the first-generation teeth.

Only the most labial (and thus oldest) first-generation teeth are preserved complete. More lingual first-generation teeth have been partly resorbed so that only their bases remain, and these bases overlap considerably (Figure 6F and 7B). In this zone, each first-generation tooth has thus been resorbed before the next, more lingual, tooth was added to the file. Albeit not basal, the resorption surfaces are wide open, not only to the next added tooth, but also to the replacement tooth bud that formed immediately above them (Figure 6C-F). In the same way as those of the tooth cushions (the primitive form of inner dental arcades; Chen et al., 2017), the first-generation teeth thus establish the tooth positions for the cyclic replacement teeth. The addition of new first-generation teeth to the lingual ends of tooth files, and the deposition of replacement teeth onto the labial tooth sockets of these files, are likely parallel processes. The former process constructs tooth files from the positions set up by the founder ridges via semi-basal resorption (Figure 6A), and the latter builds replacement columns (RT) at each position along the files via basal resorption (Figure 6B). As the successive tooth positions in a file increasingly overlap, at a point, semi-basal resorption becomes completely basal. The successive teeth in a file now take on the appearance of a replacement column, representing a marginal position that is drifting and tilting lingually (Figure 6, RC4-8; Animation 1). Intriguingly, where space allows, a new position can be inserted into the marginal replacement column like a branch (Figure 3, TR4-10 to RC4-8, TR3-10 to TR3-9). This can substitute the labial positions that have been overgrown by ornament (see below).

**Figure 7.**
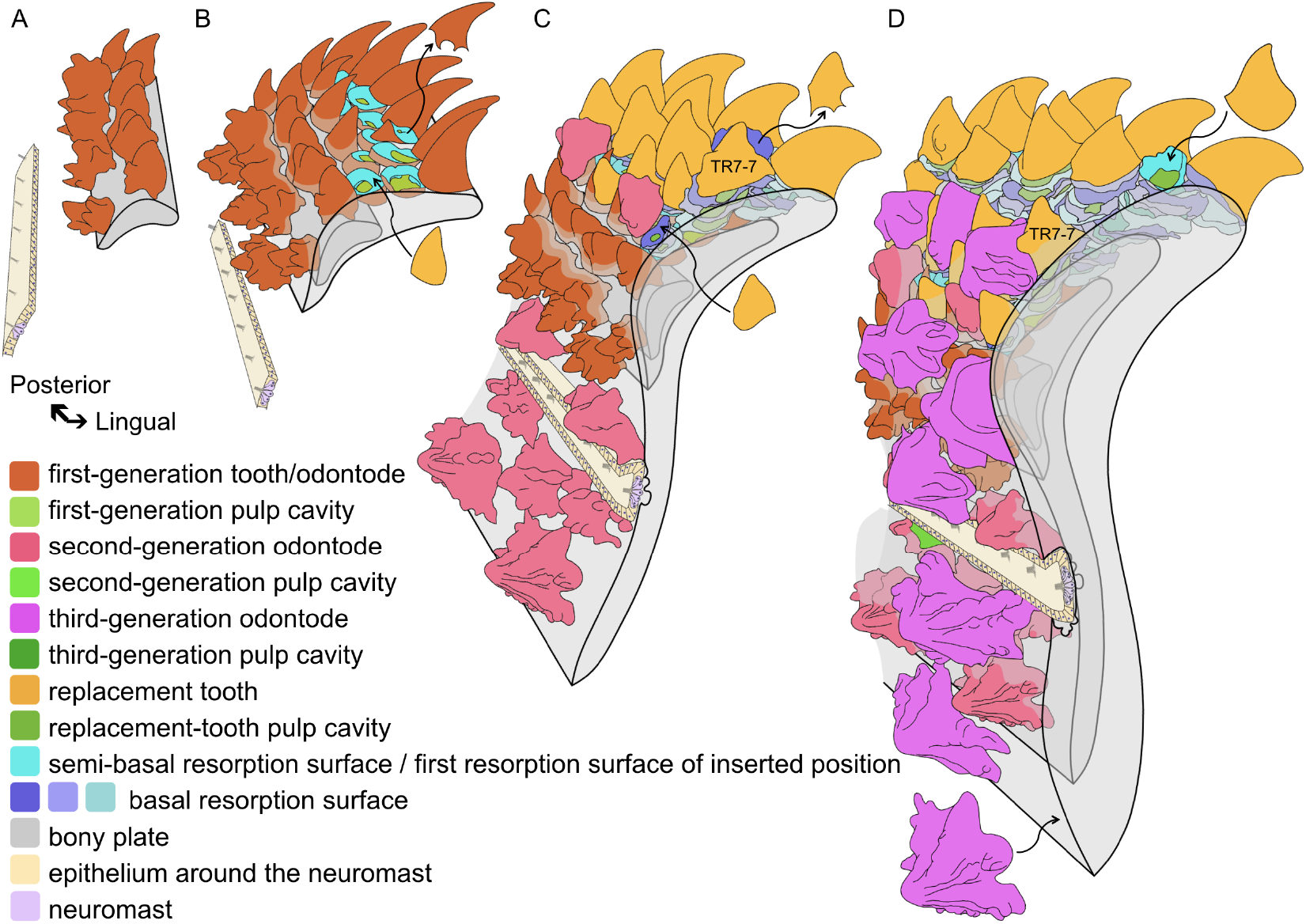
Reconstructed ontogeny of the *Lophosteus* marginal jawbone relative to development of the lateral line. Block diagrams in antero-external view, considering the rotation of bone in life. The founder ridges and the ornament-like tooth TR7-7 mark the labial rotation and drift of the tooth rows. A, the initial odontodes are formed as founder ridges. B, isolated ornament odontodes and teeth are added sequentially in opposite direction, attaining a stellate and conical morphology, respectively. The teeth are shed semi-basally, establishing replacement tooth positions. C, teeth are cyclically replaced at the positions set up by the first-generation teeth; the second-generation ornament odontodes start to invade the oral domain lingually and form around the lateral line canal labially. Note that the replacement of teeth and the overgrowth of odontodes may have started before the addition of the first-generation teeth is finished. D, the labial extension of lateral line canal partially resorbs the second-generation ornament odontodes at its labial border; the third-generation ornament odontodes overgrow more labial tooth rows that have been rotated to the face; new tooth positions are inserted to compensate the embedded labial tooth positions. Curved arrows, examples of the adding or shedding of teeth or ornament odontodes. Note that the buried part of teeth and odontodes, the embedded resorption surfaces and bone mineralized at earlier stages become increasingly grey. The lateral line is represented by neuromasts and epithelial cells. The size and number of neuromasts is schematic and only to represent their presence.

The inserted position is aligned in a tooth file with the preexisting positions, as if all the positions had been added sequentially. The actual pattern is thus more interesting and complex than it appears superficially. The most labial position has the shortest history (TF5-3 has not been replaced); the marginal position has the longest history (e.g. RC4-8 has been replaced 17 times), while the inserted position immediately labial to it has the most recent history (TR4-10 has been replaced once). Therefore, each position has a different replacement history, which can be revealed by the shape of the pulp cavity (Figure 6B and 8). Compared to the developmental history of the bone, the labial positions function at the earlier stages, depending on how soon they are overgrown by ornament, and thus the final replacement teeth of the labial tooth rows are smaller than those of the lingual ones (Figure 4B and 8, gold).

**Figure 8.**
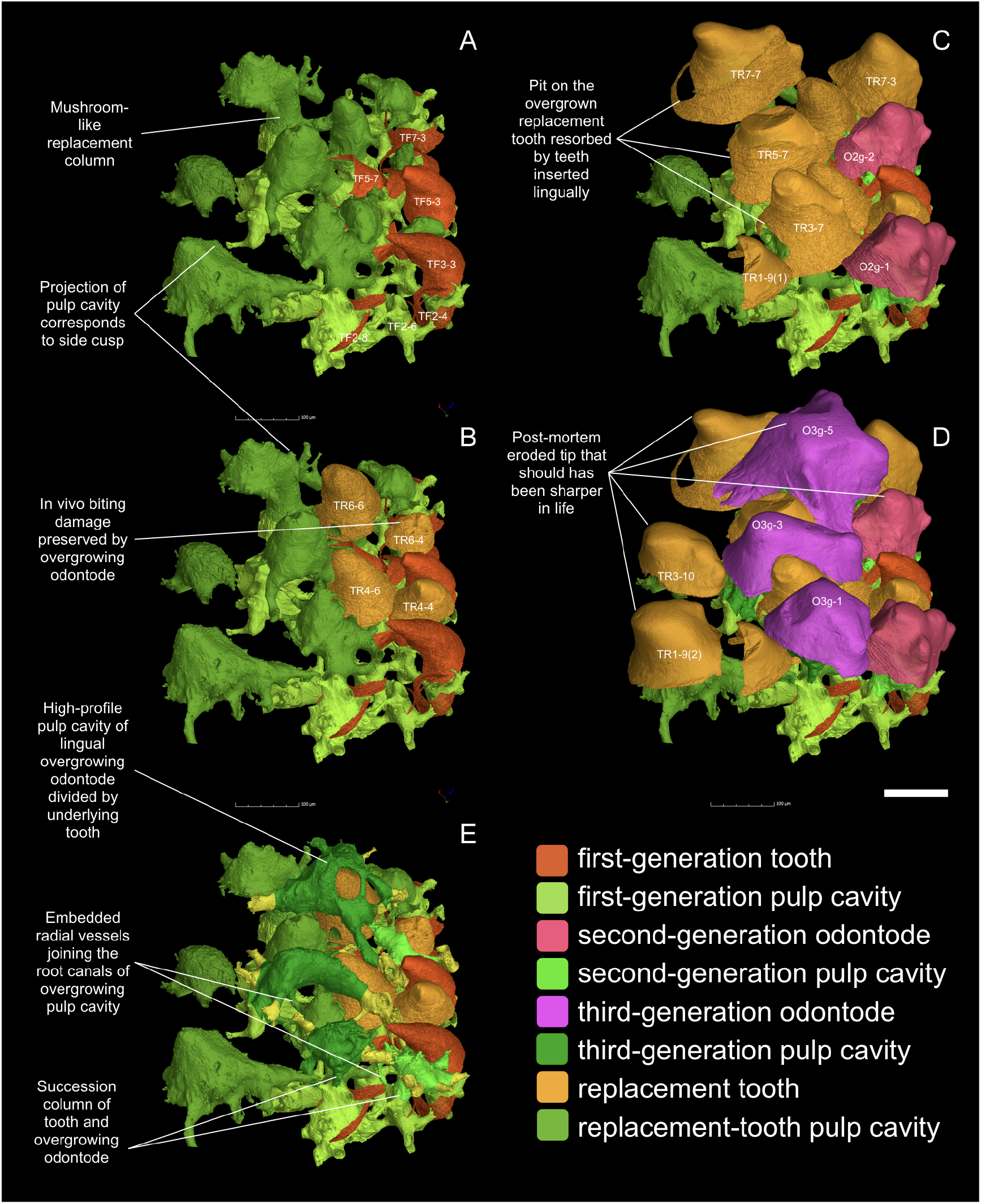
Postero-occlusal view of tooth field invaded by overgrowing odontodes. A, first-generation teeth are shown. They are perfectly conical. B, the final replacement teeth that will be buried by second-generation overgrowing odontodes are shown. They are still perfectly conical. C. the final replacement teeth that will be buried by third-generation overgrowing odontodes are shown. They are more or less ornament-like, probably because of the approach of the dermal epithelium, which is represented by the second-generation overgrowing odontodes, during the tooth development. D, the final replacement teeth of the inserted positions are shown. They are ornament-like, probably because of the approach of the dermal epithelium that generate the third-generation overgrowing odontodes. E, spatial relationship between the pulp cavities of overgrowing odontodes and teeth buried right below. Scale bar, 0.1mm.

The pulp cavities of the first-generation teeth lie directly on the basal compact bone, coinciding with the territory of the basal feeder vessels (Figure 6A). This indicates that the most lingual row of first-generation teeth was once located at the jaw margin and that the oral lamina at this early developmental stage was much narrower (Figure 3C, dashed curve; Figure 7B). Only a few of the first-generation teeth are directly supplied by the basal feeder vessels, but all of them have radiating canals to adjoin each other, forming a network on the dentine-bone interface (Figure 6A; Figure 8A, pulp cavities of TF2-6 and TF2-8). Each pulp cavities of the replacement teeth is directly fed by those of the first-generation odontodes. A first-generation tooth and all its replacement teeth have their pulp cavities fused like a mushroom (Figure 3B, RC4-4, RC4-6; Figure 8). The mushroom stem, which is the accumulated pulp cavities of the successive generations of shed teeth, indicating the directional migration of the tooth position. It becomes a long column at the lingual positions that have been replaced many times, extending beyond the area supplied by the basal feeder vessels. Hence, a horizontal vascular mesh is laid down above the compact bone of the new portion of the oral lamina (Figure 5 and 6A-B, pink), connected to a few new feeder vessels that are incorporated at the new bone margin. Vertical canals of different sizes and shapes are sprouted from the pulp column down towards the pulp column of other positions, as well as towards the horizontal vascular mesh (Figure 6B), like the adventitious prop roots of banyan trees supporting the trunk that spreads laterally.

### The Ornament

The labial founder ridge has a robust tube-like pulp cavity (Figure 5, FRla), while almost all the odontodes on the facial lamina have pillow-like pulp cavities that extend laterally beyond the crown (Figure 3, O3g-3-6, O2g-2-5). The first row of isolated spiny ornament odontodes (Figure 5 and 6A, OP0-1, OP2-1, OP4-1, OP6-1), however, only has narrow pulp canals that are much smaller than the crown (Slideshow 1B). The pulp canals are situated at the same level as all the first-generation pulp cavities, just above the basal feeder vessels, showing that these odontodes, with a similar spiny appearance as the labial founder ridge, are laid down directly on the bony plate, rather than being inserted between preexisting odontodes. They overlap the labial edge of the labial founder ridge (Figure 3, O1g-2-1), like the younger first-generation teeth overlapping the lingual edges of the older ones on the oral lamina. All this suggests that they are added immediately after the labial founder ridge, at the labial margin of the freshly mineralized jawbone. The single row of spiny odontodes, the three rows of the two founder ridges and the several rows of first-generation teeth together comprise the first generation of odontodes, corresponding to the jawbone of the early stage (Figure 3, dashed curve; Figure 4B, orange; Figure 7A-B). As the facial lamina extends labially, younger generations of odontodes are initiated to cover the new bone (Figure 7C-D).

Unlike the teeth, no resorption occurs in the dermal odontodes. Younger generations of larger odontodes thus simply overgrow, rather than replace, the preexisting dermal odontodes. There are two levels of overgrowth and the dermal odontodes are divided into three generations, judging from their size and distribution (Figure 5). The dermal odontodes have their pulp cavities squeezed if they are deposited on top of a cusp (Figure 3, O3g-4, O3g-7; Figure 8E).

An odontode can be altered by encountering the lateral line canal. The second-generation odontodes situated along the lingual border of the lateral line canal have their labial ridgelets truncated, in order not to intrude on the canal sulcus (Figure 4B, O2g-3-2, O2g-7-2). The alternate positions that are supposed to form the next row are suppressed (Figure 4C, null sign). This indicates that the lateral line exists in the epithelium before being canalized by the jawbone (Figure 7A-B), and signals to the odontodes not to encroach on it. But this does not stop the odontodes from spreading across the lateral line canal with the same pattern: alternate positions can develop properly on the other side of the canal (Figure 5B, OP1-4, OP5-4). When the lateral line canal grows wider by extending labially, the lingual edge of the odontodes at the labial border is resorbed, making their pulp cavities open to the canal (Figure 3, O2g-3-3; Figure 4A, asterisk; Figure 5B, OP3-3, OP7-3; Figure 7D; Slideshow 1C). The ridgelets of the third-generation odontodes overgrowing on both borders of the canal are also compressed (Figure 4A, O3g-4, O3g-7). Like those supporting the odontodes, the feeder vessels enter the bottom of the lateral line canal in a labially oblique direction (Figure 2J_2_ and 3). But all those supporting the odontodes form primary osteons with a cementing surface, in sharp contrast, the feeder vessels of the lateral line canal are wrapped by irregular resorption surfaces, forming secondary osteons. It indicates that bone remodeling has occurred in the lateral line region (Figure 2B and 7C-D; Slideshow 1E).

Contrary to the lingual jaw margin that is packed with replacement columns, the labial margin is not fully covered by odontodes (Figure 4A). On the uncovered labial margin, small vessels are incorporated in the upper layer of the bone (Figure 5A, yellow vessels with dashed arrows). They probably originate from the skin vascular system and extend lingually to join the radial vessels of the previous pulp cavities, sharing the openings that are arranged in a ring surrounding each odontode. They would have turned into the radial vessels that emerge from the pulp cavities of the unattached odontodes, when the odontodes were fixed onto the bare labial margin.

The small vessels leading to the putative developing odontodes (Figure 5, OP6-7) at the labial margin are connected to some of the second most labial odontodes (OP6-5), instead of the most labial ones (OP3-6). This is in accord with the alternate organization of all the isolated odontodes that are laid down directly on the basal bone. The labial founder ridge does not have the main cusps divided into two alternate rows like the lingual founder ridge, but possesses both odd- and even-number positions (Figure 4B-C). Although the first-generation spiny odontodes and the first-generation teeth are generated in the opposite directions from the founder ridges, they set up the same number of odontode files and tooth files. If the first row of the first-generation teeth establishes the odd-number tooth files, then the second row initiates the even-number tooth files. Each of the more labial rows of stellate odontodes are also deposited between two odontodes of the previous row. This pattern persists from generation to generation, and has been carried on across the lateral line canal (Figure 5B). However, the increase in size of the stellate odontodes through generations is more dramatic than the growth in length of the jawbone. Consequently, not all the gaps between odontodes of the previous row are filled by new odontodes, and the number of stellate odontodes in a row decreases labially. The first-generation odontode row (Row 1) crosses all the even-number files, taking every second positions (0, 2, 4, 6); but the second-generation odontodes only occupy every second odd-number positions (1, 5 and 3, 7) in Row 2-4 and every second even-number positions (2, 6) in Row 5, and the third-generation odontodes occupy every third odd-number positions (3, 9) in Row 6. Accordingly, we can predict that the next odontode of Row 7 (OP6-7)will be added in File 6 to fill the gap between OP3-6 and OP9-6.

Younger generations of stellate odontodes not only extend further down the facial lamina but also begin to invade the oral lamina (Figure 5, O2g, O3g). Their pulp cavities tend to fill the gaps between rows of preexisting odontodes (Figure 3, O3g-2, O3g-3; Figure 8E; Slideshow 1B-C, O3g-7). The shape of the pulp cavities is not necessarily consistent with the overgrowing odontodes, for instance, O3g-2 deposited right on top of the lingual founder ridge has its pulp cavity divided into two parts to fit the gaps on either side of the ridge (Figure 4A, 5A and 6B).

As mentioned above, side cusps are most abundant on the labial flank of the labial founder ridge, while the main cusp of isolated odontodes in the same row is surrounded by sharp side cusps. In the next row, the first-generation odontodes are longitudinally elongated, with the side cusps tending to be aligned radially. The still more labial odontodes of younger generations gradually attain the typical stellate morphology by narrowing the side cusps into ridgelets (Figure 4B). As in the teeth and founder ridges, orthodentine extends all the way down to the pulp cavities in the first generation of odontodes. In the younger generations, orthodentine is limited to the periphery; the interior is filled by denteons (dentinal osteons, Ørvig, 1951, 1967), which turn into cellular osteons at the base (Slideshow 1A-C). The bone-like tissue surrouding the lower part of the ascending canals above the pulp cavity (Figure 3C; Gross, 1969, afk; Jerve et al., 2016), as well as the root canals beneath the pulp cavity (Figure 3C; Gross, 1971b, ms), appears externally as the bone of attachment (Figure 4A, in grey). By contrast, the ornament odontodes overgrowing the oral lamina do not develop a fully stellate morphology, with few uncrenulated ridgelets. The most lingual ones tend to become longitudinally compressed, contrary to the longitudinal elongation of those on the facial lamina. Their side cusps only develop labially and main cusps incline lingually (Figure 4A and 8D, O2g-2, O3g-3). Their pulp cavities are surrounded by acellular dentine (Slideshow 1C). To put it another way, they come to look somewhat like the teeth, whose territory they are invading.

Overall, all generations of ornament odontodes, including the first one, follow a global morphological gradation. The further they are from the biting margin at the time they are added, the more side cusps, ridgelets and nodules are associated with the main cusp, and the more ascending canals are attached to the flattened pulp cavities. The largest and youngest stellate odontodes have the most nodular appearance (Figure 4A, O3g-3-6, O3g-7) and the most osteodentine-like tissue (Slideshow 1A-D), which probably reflect their location furthest from the jaw margin, rather than their size or age. We surmise that this pattern reflects a morphogenetic signal gradient (Figure 4) between the jaw margin and the “generic dermal surface” of the face, represented here by the facial lamina beyond the lateral line canal.

### Interaction between teeth and ornament

A subtle but important point of morphological variability affects the labial surface of some of the shedding teeth. Replacement teeth at the lingual jaw margin, as well as the first-generation teeth, always have a smooth labial surface (Figure 8B), but teeth in positions close to invading ornament odontodes are often nodulated on the labial side (Figure 4A and 8C-D, arrow head, TR1-9(2), TR3-7, TR3-10, TR6-9, TR7-3, TR7-7). Since the pulp cavities of teeth are not infilled as much as those of ornament odontodes, the pulp cavities of the ornament-like teeth show projections of nodules as well (Figure 8, pulp cavities of TR1-9(2) and TR7-7). The dentine of the labial nodules tends to include some cell spaces. In other words, just like the most lingual ornament odontodes display tooth-like characteristics (see above), these teeth show ornament-like characteristics; there appears to be a degree of “morphological cross-contamination” between the two odontode sets (Figure 4). At the invasive front line, both overgrowing odontodes and replacement teeth have a half-tooth half-ornament morphology (Figure 8C, compare O2g-1 and O2g-2 with TR7-3 and TR7-7). Notwithstanding the morphological similarity, the replacement teeth are instantly recognizable by their possession of basal resorption surfaces. Since the invasive front line of the ornament is irregular, the morphology of teeth is not correlated with rows or generations, but with the proximity of overgrowing odontodes.

For the tooth positions initiated by the first-generation teeth, the rule is that the more lingually they are located, the more times they have been replaced. This seems not caused by a higher replacement rate of the lingual positions, but the overgrowth of ornament on the labial positions. When a tooth position is buried by ornament, even just partially, its replacement cycle will be terminated, probably due to the blocking of the radial vessels causing the failure of basal resorption (see Chen et al., 2017). Its final replacement tooth cannot be shed, but might be slightly encroached by the basal resorption of the tooth positions lingual to it (Figure 8C, TR1-9(1), TR3-7, TR5-7, TR7-7). Some first-generation teeth have been resorbed semi-basally but cannot set up a cyclically replaced position because their open pulp cavities are quickly taken by overgrowing odontodes. Put differently, these first-generation teeth are replaced by a tooth-like dermal odontode (Figure 6C-D and 8, TF2-6 replaced by O2g-1).

## Discussion

### Comparative context: the dentitions of *Andreolepis* and ‘acanthothoracids’

The marginal dentitions of the stem osteichthyan *Andreolepis* (Late Silurian, Gotland, Sweden) and the ‘acanthothoracid’ stem gnathostomes *Kosoraspis* (Early Devonian, Prague Basin, Czech Republic) are of particular interest as comparators for *Lophosteus* because they show a similar combination of transverse alternate tooth files with lingual tooth addition, carried on marginal bones that also bear dermal ornament odontodes on their deep facial laminae (Chen et al., 2016; Vaškaninová et al., 2020). *Lophosteus* aligns with *Andreolepis* and differs from stem gnathostomes in showing resorptive tooth shedding, a unique osteichthyan characteristic (Chen et al., 2016). Indeed, this process is of fundamental importance in shaping the dentition of *Lophosteus*, with some tooth positions showing as many as twenty resorption-replacement cycles. This has important implications for understanding the ontogenetic history underlying the final adult morphology.

*Kosoraspis* has a much simpler ontogenetic history without resorption-replacement cycles, essentially corresponding only to the first-generation odontodes of *Andreolepis*. *Kosoraspis* show a unidirectional change from small ornament odontodes at the labial margin of the jawbone, through gradually larger and progressively more tooth-like ornament odontodes, to teeth (Vaškaninová et al. 2020). It indicates the ossification center locates at the labial margin. The gradient between teeth and dermal odontodes of the first generation appears to be unidirectional in *Andreolepis* too, from a morphology with elongate bases to a more obviously tooth-like morphology with round bases as they approach the jaw margin (Chen et al., 2016, fig. 2a,b). Any equivalent of the founder ridges of *Lophosteus*, if any, have not been captured by the high-resolution scan that only covers approximately a quarter of the height of the facial lamina (Chen et al., 2016, fig. 1b). Nevertheless, it is certain that the location of the odontode founder region and the bone ossification center is considerably labial to the original oral-dermal boundary in *Andreolepis*. Whereas, the three developmental landmarks overlap in *Lophosteus*, which thus displays a bidirectional morphological gradation in the initial odontode skeleton.

### Developmental implications: Initiation of the odontode skeleton

The dermal odontodes and teeth of *Lophosteus* are initiated simultaneously in the form of two parallel and closely spaced founder ridges (Figure 4B; Figure 7A-B). The cusps of the lingual founder ridge incline lingually, and those of the labial founder ridge, labially; subsequent odontodes are added to the lingual flank of the lingual founder ridge and the labial flank of the labial founder ridge. This geometrical layout strongly suggests that the initiation site for odontode formation is the boundary between two patterning domains: a labial domain where the odontodes become ornament and a lingual domain where they become teeth. Because the development of an odontode is always initiated by an epithelium, we infer that the two domains are covered with two distinct epithelia, which we will refer to respectively as the dermal and oral epithelium. We have no direct evidence for their cellular or molecular identities, but one obvious possibility is that the dermal epithelium is ectodermal whereas the oral epithelium is at least partly endodermal. Indirect support for this hypothesis is provided by lineage tracing in developing sturgeon (Minarik et al., 2017), which reveals an extensive endodermal epithelial contribution to the jaw margins. The cusps of the two founder ridges are already morphologically distinct, and the odontodes that are subsequently produced at their labial and lingual sides quickly acquire the full characteristics of ornament and teeth, respectively. This must represent the establishment of their complete regulatory cascades, and possibly relates to the initiation of new odontodes on each side moving away from the boundary between the dermal and oral epithelia into the presumably more homogeneous signaling environment of a single epithelium (Figure 4, 1g).

### Developmental implications: the shifting oral-dermal boundary

The strict, linear separation described above only characterises the first-generation odontodes. Already in the second generation, stellate odontodes have overgrown some of the oldest teeth, in a considerably more lingual position than the original boundary. The third generation penetrates even further into the territory of teeth (Figure 4, 2g, 3g; Figure 5, O2g, O3g). This suggests an irregular expansion of the dermal epithelium into the region previously occupied exclusively by the oral epithelium. A similar invasion of the tooth field by dermal ornament is observed in *Andreolepis* (Chen et al., 2016) and an unnamed acanthothoracid from the Canadian Arctic, specimen CPW.9 (Vaškaninová et al., 2020).

In *Lophosteus*, the ornament invasion produces an effect that is highly informative about the relationship between these two odontode sets. Essentially, the *deposition* of odontodes remains characteristic for the two sets – teeth continue to be replaced cyclically until overgrown by ornament, and the ornament odontodes are never shed – but the *morphology* of each set seems to become influenced by the other. The teeth nearest to the ornament bear side cusps or nodules that tend to form a labial ridgelet, as well as a conical main cusp which points lingually. Conversely, the most lingual ornament odontodes are small, with short side ridges that are restricted to the labial side of the odontode and carry few if any nodules (Figure 4A-B and 8C-D). That is to say, in the invasion zone, the teeth are ornament-like and the ornament odontodes are tooth-like. The simplest explanation for this phenomenon is that the invasion zone provides a mixed set of morphogenetic signals, because it is a patchwork of dermal and oral epithelium, and that both ornament odontodes and teeth respond to both signals. Note, however, that this “regulatory cross-contamination” only affects the morphology, not the deposition and (if present) resorption cycles.

In *Andreolepis*, the lingualmost odontodes of any generation of invading ornament can be tall and bear biting chips, in contrast to the characteristic flat-topped ornament morphology (Chen et al., 2016, Extended Data Figure 3c). In many basal actinopterygians with acrodin-capped teeth, such as *Birgeria* and *Boreosomus,* the dermal odontodes labial to the jaw teeth also bear an apical wart of acrodin (Ørvig, 1978a, PP. 38, Figs 3–4, 7–9). The zone of the acrodin-bearing odontodes varies in size, being, for example, narrower in *Colobodus* and wider in *Nephrotus* and others (Ørvig, 1978c, pp. 307), but is invariably restricted to the vicinity of dentition, regardless of the type of jawbone (Ørvig, 1978a, pp. 41-42). Similar phenomena can be seen in stem chondrichthyans. The labial face of the blade-like teeth on the tooth whorls of *Ptomacanthus* can be ornamented, like the tesserae that they are interlocked with (Miles, 1973). The labio-lingual rows of extraoral denticles on the whorl-like cheek scales, pointed lip scales, platelets or tesserae labial to the jaw margin of ischnacanthid acanthodians (Gross, 1971a, Tafel 4, Fig. 24-29; Ørvig, 1973, Text-fig. 1C; Blais et al., 2011, 2015; Burrow et al., 2018) also suggests the presence of a mixed dermal-oral signaling environment extending beyond the mouth. Together with these observations, the tooth-like ornament and the ornament-like teeth of *Lophosteus* call into question the demarcation between dermal and oral developmental domains.

The description of ornament invasion presented above incorporates the assumption that the position and orientation of the jawbone is static relative to the edge of the mouth. In fact, the bone appears to have rotated labially during growth (Figure 7), so that the location of the mouth margin shifted during ontogeny from the lingual founder ridge, via each lingual row of the first-generation teeth, to each successive row of replacement teeth at the current lingual margin of the oral lamina. This implies that the oral-dermal epithelial boundary do not drift lingually to any great degree; rather, the rotation of the bone causes the labial tooth rows to move onto the face, where they get covered by dermal epithelium and overgrown by ornament. This is comparable to the rotation of the tooth whorls at the jaw margin of primitive chondrichthyans, with the post-functional teeth slid beneath the skin (Smith and Coates, 2001, Figure 14.3(a); Williams, 2001).

### Developmental implications: alternate pattern throughout oral-dermal domains

In *Lophosteus*, the first-generation teeth are added sequentially towards the growing lingual margin, with the tooth families arranged in horizontal alternate files. In the next stage, each successor, if not overgrown by invasive odontodes, turns into an initiator and sets up its own tooth family by cyclic replacement. The replacement teeth hence inherit the pattern of the first-generation teeth, generation after generation, forming vertical alternate columns. The replacement columns, including those inserted later and those disturbed by overgrowing odontodes, are aligned transversely parallel to the files of the first-generation teeth (Figure 6 and 8). Longitudinally, the exposed tooth rows appear somewhat irregular, which is due to two reasons. Firstly, new positions that are not established by the first-generation teeth are inserted whenever space is available along the marginal replacement column (e.g. Figure 6B, RC3-10). Secondly, different labial positions are terminated randomly by different generations of overgrowing odontodes, and the total number of replacement cycles is different in each tooth position. As a result, the final replacement teeth drifted lingually from their first-generation teeth for a variable distance, reflected by the variable length of replacement columns (Figure 8, compare TR7-3 with TR 6-4 and TR4-4). Nevertheless, the marginal positions that maintain the original pattern throughout the growth of bone are invariably aligned in a row.

Sequential addition along the files established by the first generation also applies to the stellate odontodes on the facial lamina. Such an alternate arrangement is constant throughout the teeth and tooth-like odontodes on the marginal jawbone of *Andreolepis* and *Kosoraspis* as well, irrespective of whether they will be overgrown by younger generations of odontodes (Chen et al., 2016; Vaškaninová et al., 2020). The alternate pattern of teeth and ornament of *Lophosteus* is already established by the founder ridges. The ornament differs from teeth in the fact that every other alternate position of ornament is suppressed as the ornament extends labially (Figure 5B); this is because consecutive generations of ornament odontodes increase in length more quickly than the bone. The replacement of two teeth by a single larger successor at a crowded site has been observed in frogs and lizards, and considered as a common phenomenon (Gillette, 1955; Edmund, 1960; Cooper, 1963; Osborn, 1971). The suppression does not represent an irregularity; instead, it may reflect the fundamental mechanism producing the alternate pattern.

The hexagonal pattern is the most efficient form of close packing of rounded objects. The close packed teeth of myliobatid rays, which have turned into short hexagonal prisms (Edmund, 1960; Underwood et al., 2015), is an extreme example of a dentition imitating the structure of a honeycomb. A regular pattern, which was thought to be unique to teeth and reflect the spatio-temporal regulation of the dental lamina (Smith, 2003; Underwood et al., 2016), is actually not uncommon in dermal odontodes, as well as in bony denticles. Ordered tubercles can be seen covering the armor of jawed stem gnathostomes as long as there is only one generation, irrespective of whether they are dentinal units on the gnathal plates and spines, or bony units on the postbranchial lamina (Bystrow, 1957, Fig. 2; Burrow, 2003; Johanson and Smith, 2003, 2005; Young, 2003).

An alternate pattern can be produced by odontodes that are laid down directly on the bony plate, simply through filling the gaps between the odontodes in the previous row, even if the previous row is from the older generation (Figure 5B). But it is difficult for the enlarged successive odontodes to find and fit such gaps, by reason of the preexisting odontodes, so the pattern will be disturbed. This effect is clearly visible in a digitally dissected spine of the jawed stem gnathostome *Romundina,* which carries three generations of odontodes (Jerve et al., 2017, fig. 2D1-D3); the first generation shows an ordered pattern, but this breaks down in the second and third generations. The single-file arrangement of the first-generation odontodes of *Andreolepis* scale is also obscured by the overgrowing odontodes (Qu et al., 2013).

All these examples agree with a fundamental embryologic mechanism of odontode patterning shared among the skeletons of vertebrates, which had evolved prior to the origin of teeth, as already proposed by Osborn (1971). The alternate pattern can be self-generating, as long as the size of inhibitory zones is equivalent or in a smooth gradient (Osborn, 1977). Therefore, the regularity of organization should not be considered as a criterion of true teeth. The claim that the ectoderm (dermal epithelium) lacks patterning capacity (Fraser and Smith, 2011; Smith and Johanson, 2015), which has been used to support the idea of a fundamental difference between dermal odontodes and teeth, is biased by the derived adult condition of modern chondrichthyans. Actually, sharks also have regular rows of skin denticles at an early developmental stage, including the caudal denticles, dorsal denticles and the first-generation general denticles (Martin et al., 2016). Later on this spacing is obscured, probably by the accidental shedding of original scales and the replacement by new scales of variable sizes, at different developmental stages. Wound-healing experiments on sharks show that the loss of the diagonal rows and the rostro-caudal polarity as well as the regular size and shape of the scales in repaired squamations is caused by disturbance of the diagonal arrangement of the anchoring collagen fibers (Reif, 1978). By contrast, the regular and orderly shedding and replacement of chondrichthyan teeth preserves the embryonic pattern.

### Evolutionary developmental relationship between teeth and ornament

Our data from *Lophosteus* are in two respects uniquely informative about the relationship between teeth and ornament. Firstly, as a Silurian stem osteichthyan, *Lophosteus* represents a very short phylogenetic branch in a basal part of the gnathostome crown group, and is thus likely to present primitive characters for the Osteichthyes and maybe for the Gnathostomata as a whole; secondly, we can trace the developing relationship between teeth and dermal odontodes through the life history of the animal, whereas all the dermal odontodes and teeth that have been compared in previous studies are fully differentiated forms in adults.

The unified arrangement of the teeth and ornament of *Lophosteus* challenges the currently popular idea that teeth and dermal denticles have fundamentally different patterning regimes (Fraser and Smith, 2011). In *Lophosteus*, teeth and ornament are never starkly different. The cyclic replacement teeth can have an ornament-like appearance, and can be partially shed and overlapped. The ornament odontodes can be added sequentially and alternately, and organized in rows and files. New tooth positions can be inserted once there is a gap, breaking the original addition sequence.

Looking further afield, it is noteworthy that the “extra-oral teeth” that occur in some teleosts also display a regular organization, development in epithelial invaginations, and shedding and replacement by basal resorption of the attachment bone and supporting bone (Sire and Huysseune, 1996; Sire et al., 1998; Sire, 2001; Sire and Allizard, 2001). These structures reveal the potential plasticity of odontodes and suggest the conventional criteria of “true teeth” are not in fact unique to oral teeth.

We propose, on the basis of these data, that teeth and dermal odontodes are in essence modifications of a single odontode system. The fossil evidence from *Lophosteus* and *Andreolepis* shows that, within the ontogenetic trajectory of an individual, there could be signal cross-communication between the dermal and oral domains, leading to hybrid odontode morphologies. Interestingly, the primordial ornament odontodes of the scales of early osteichthyans, such as the ganoid scales of *Andreolepis* or the cosmoid scales of *Psarolepis* and *Porolepis* (Bystrow, 1959; Qu et al., 2013, 2016), have a more pointed and tooth-like morphology than the overgrowing odontodes. This resembles the relationship between the founder ridges and ornament odontodes of the jawbone of *Lophosteus*, suggesting that the earliest odontodes may be similar all over the body, with later deposited odontodes differentiating according to their location and function. The earliest developmental stage of the odontode skeleton may have been lost in derived taxa or, if present, missed by the convectional investigative techniques. Direct comparison between the substantially modified subsets of odontodes, could lead to the claim that teeth and dermal odontodes are two wholly separate systems. Crucially, *Lophosteus* shows us that only ontogenetic data going back to the earliest stages of development are able to reveal the original patterning relationships within the odontode skeleton.

## Material and Methods

### Specimens collecting and photographing

The marginal jawbone fragments of *Lophosteus* were collected from fallen blocks of limestone at Ohesaare cliff, Saaremaa Island, Estonia, the type locality of *Lophosteus.* Acetic acid dissolution and extraction of the microremains were carried out at the Department of Earth Sciences of Lund University and the Department of Organismal Biology of Uppsala University, Sweden. The specimens are registered to the Geological Institute, Tallinn, Estonia GIT 760-12~28. All specimens in Figure 1 were photographed under an Olympus SZX10 microscope and camera setup, including ImageView imaging software, at the Swedish Museum of Natural History, Stockholm.

### PPC-SRμCT

The specimen was imaged at beamline ID19 of the European Synchrotron Radiation Facility (ESRF) in Grenoble, France, using propagation phase-contrast synchrotron radiation microtomography (PPC-SRμCT) adapted to fossil mineralized tissues histology (Tafforeau and Smith, 2008; Sanchez et al., 2012). With an isotropic voxel size of 0.696μm, the scan was obtained with an objective 10×, NA0.3 coupled with a 2× eyepiece. The optics, associated with a gadolinium gallium garnet crystal of 10μm thickness (GGG10) scintillator, is coupled to a FreLoN 2K14 detector (fast readout low noise camera; Labiche et al., 2007) used in full frame mode with a fast shutter. The specimen was set up 15mm from the optics. The gap of the undulator U17.6 was set to 20mm and provided a pink beam (direct beam with a single main narrow harmonic) at an energy of 19keV, filtered by 0.7 mm of aluminum. A total of 4998 projections of 0.3s each were taken over 360° by half-acquisition of 600 pixels. Reconstruction was done with a modified version of a single-distance phase retrieval approach (Paganin et al., 2002; Sanchez et al., 2012).

The virtual thin sections of the sample in the form of stacks of images were segmented into three-dimensional sub-volumes through the software VG Studio 3.1. Embedded subtle structures, such as the surfaces of resorption and dentine, were traced manually.

### Odontode numbering

Each standing odontode, including ornament odontodes (O), first-generation teeth (TF) and replacement teeth (TR), is numbered according to its location and generation. Pulp cavities are numbered to denote ornament positions (OP), tooth positions (TP) and replacement columns (RC). Normally they are marked as “file number-row number”. A replacement column is formed by the succession of teeth at a tooth position. Replacement columns that derive directly from a first-generation pulp cavity have the same number as the first-generation teeth, i.e. the position number. The youngest inserted positions are not aligned in rows, but all of their row numbers are set as 10 in order to be differentiated from the others. The generation of replacement teeth varies dramatically among positions, and thus is not indicated (though it can be calculated from the number of resorption surfaces); but ornament odontodes are distinguished by generation numbers (1g/2g/3g), which is added before the file numbers. For example, O2g-6-5 represents the second-generation ornament odontode that is situated in the sixth file of the fifth labial row. File numbers can be overviewed from Figure 4C (7 files in the modeled area), and generation number are illustrated by colors. The file/row numbers are not applicable to the overgrowing odontodes that are located irregularly, and “generation number-serial number” is used instead. For instance, the “7” of O3g-7 does not related to File 7 or Row 7.

## Supporting information

Slideshow 1

Animation 1

## Supplementary files

Slideshow 1. Virtual thin sections. Sectioning planes are indicated by dashed lines on Figure 3D. Scale bar, 1 mm.

Animation 1. Tooth addition and replacement of two alternate files. Perspective view.

## Acknowledgements

We thank ESRF for granting beamtime to the scanning proposal (ES151, D.C.) and the Knut & Alice Wallenberg Foundation for funding (Wallenberg Scholarship, P.E.A.).

## Author contributions

Material collecting and sorting: T.M., H.B., P.E.A. and D.C. Conceptualization: D.C., P.E.A. and H.B. Scanning: S.S., P.T., D.C., and P.E.A. Raw data reconstruction: S.S. and P.T. 3D modeling: D.C. Technical support: S.S. Interpretation: D.C. Figure and animation preparation: D.C, except Figure 1 by P.E.A and Figure 2 by H.B. Writing: D.C. and P.E.A., with input from all other authors.

## Data accessibility

All original segmentation is available through the ESRF paleontology database.

