## Supplementary figures and images for "Dental ontogeny in the most primitive bony fish *Lophosteus* reveals the developmental relationship between teeth and dermal odontodes"

### Slideshow 1

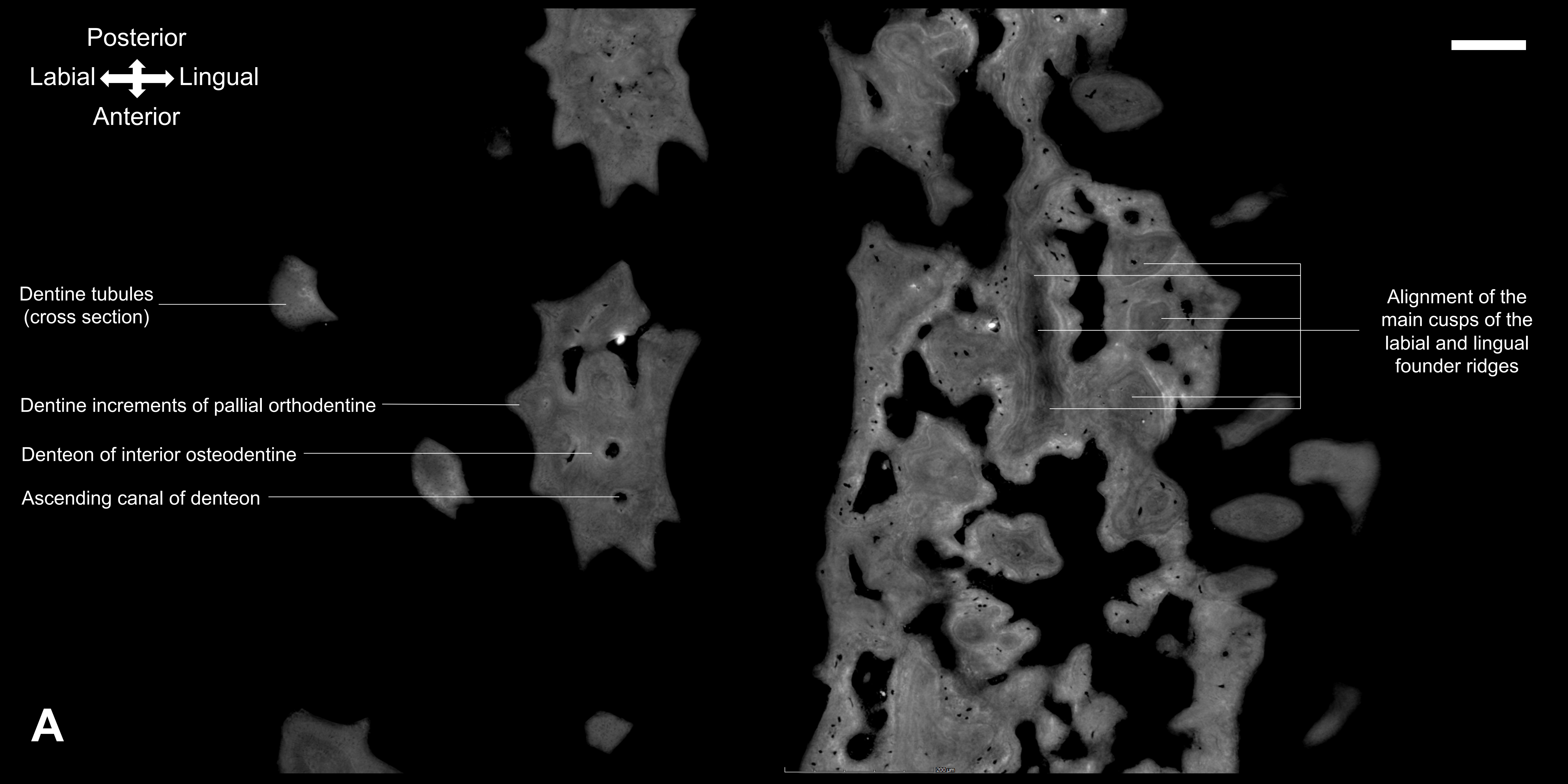

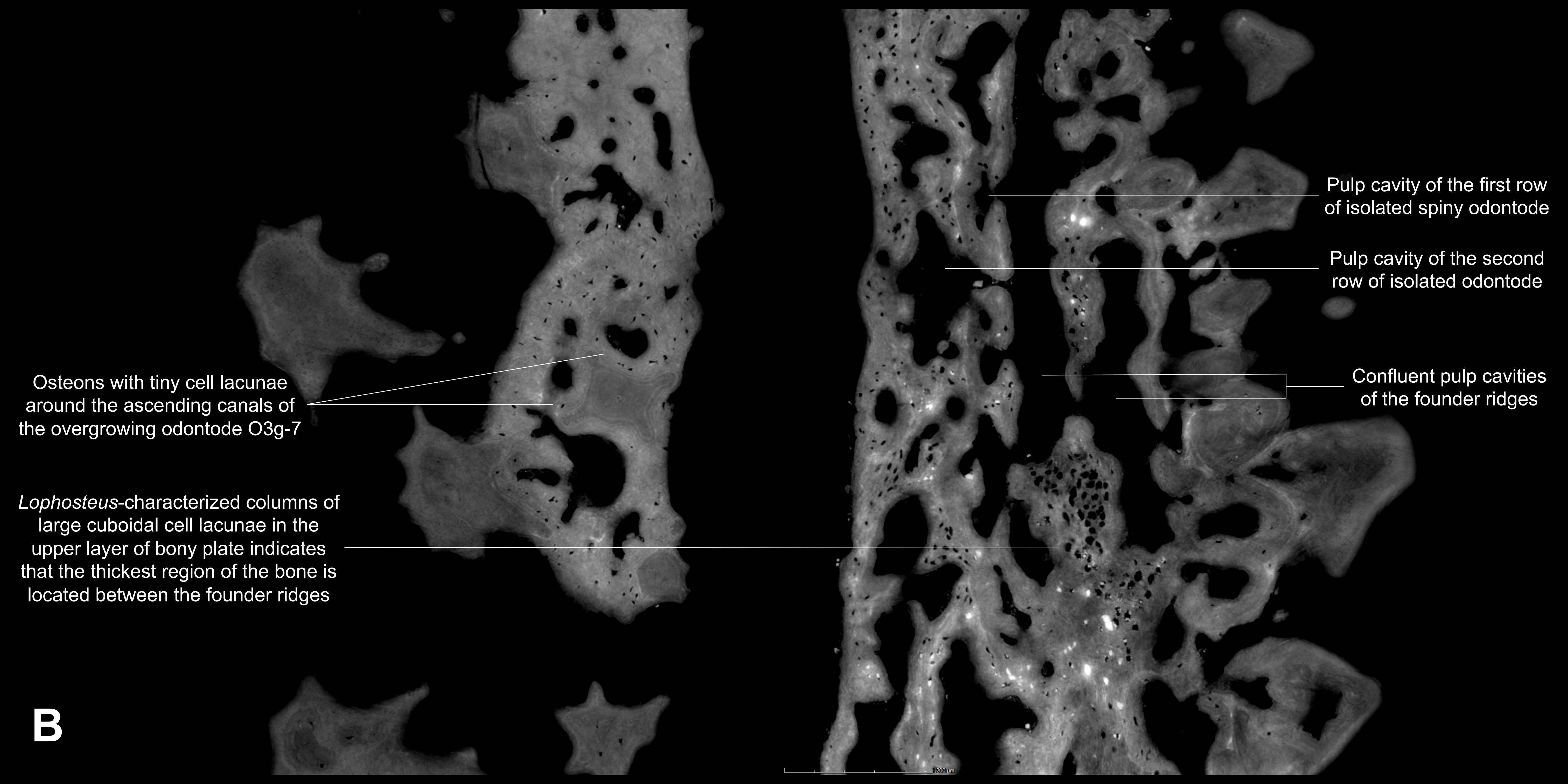

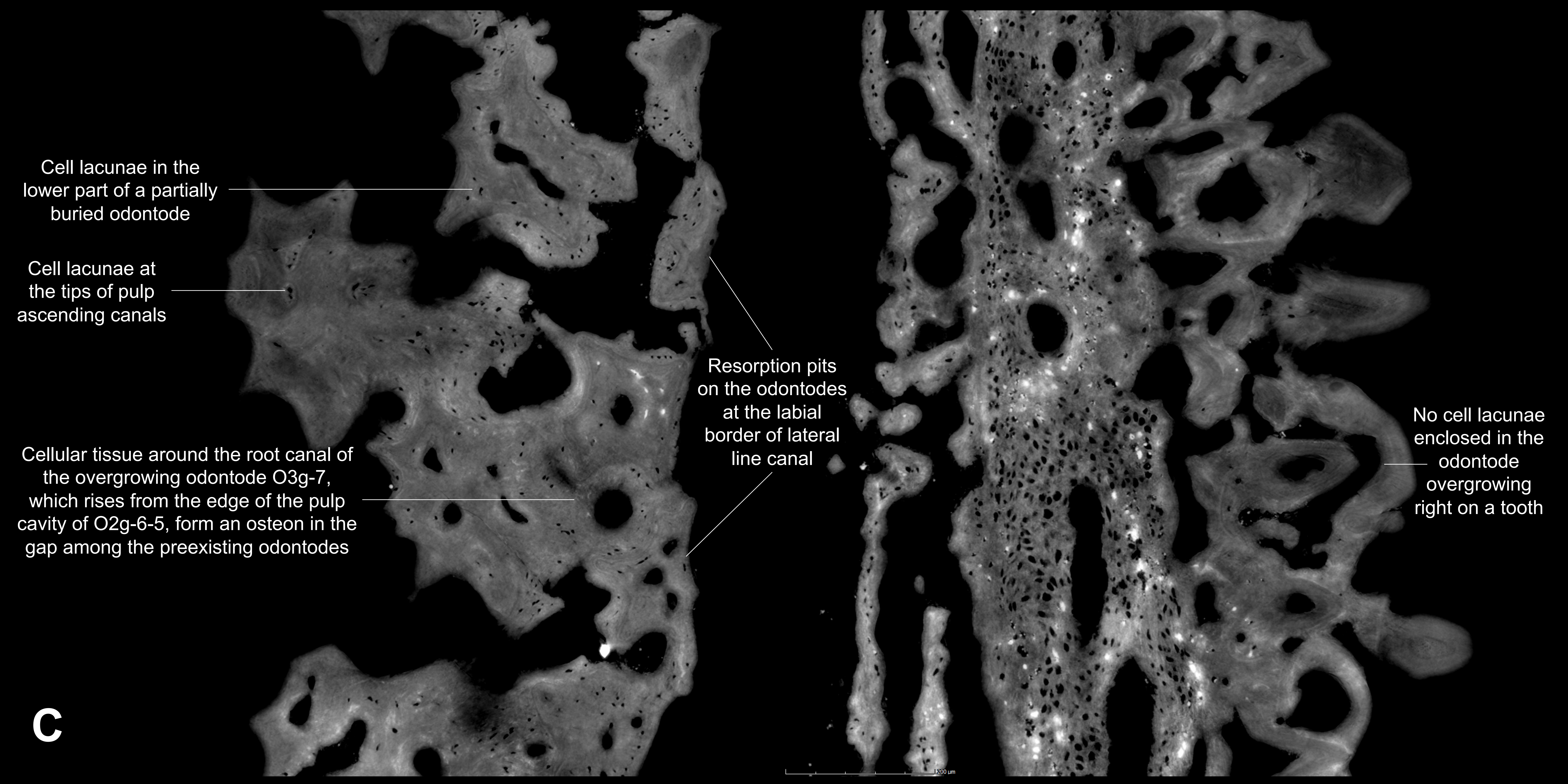

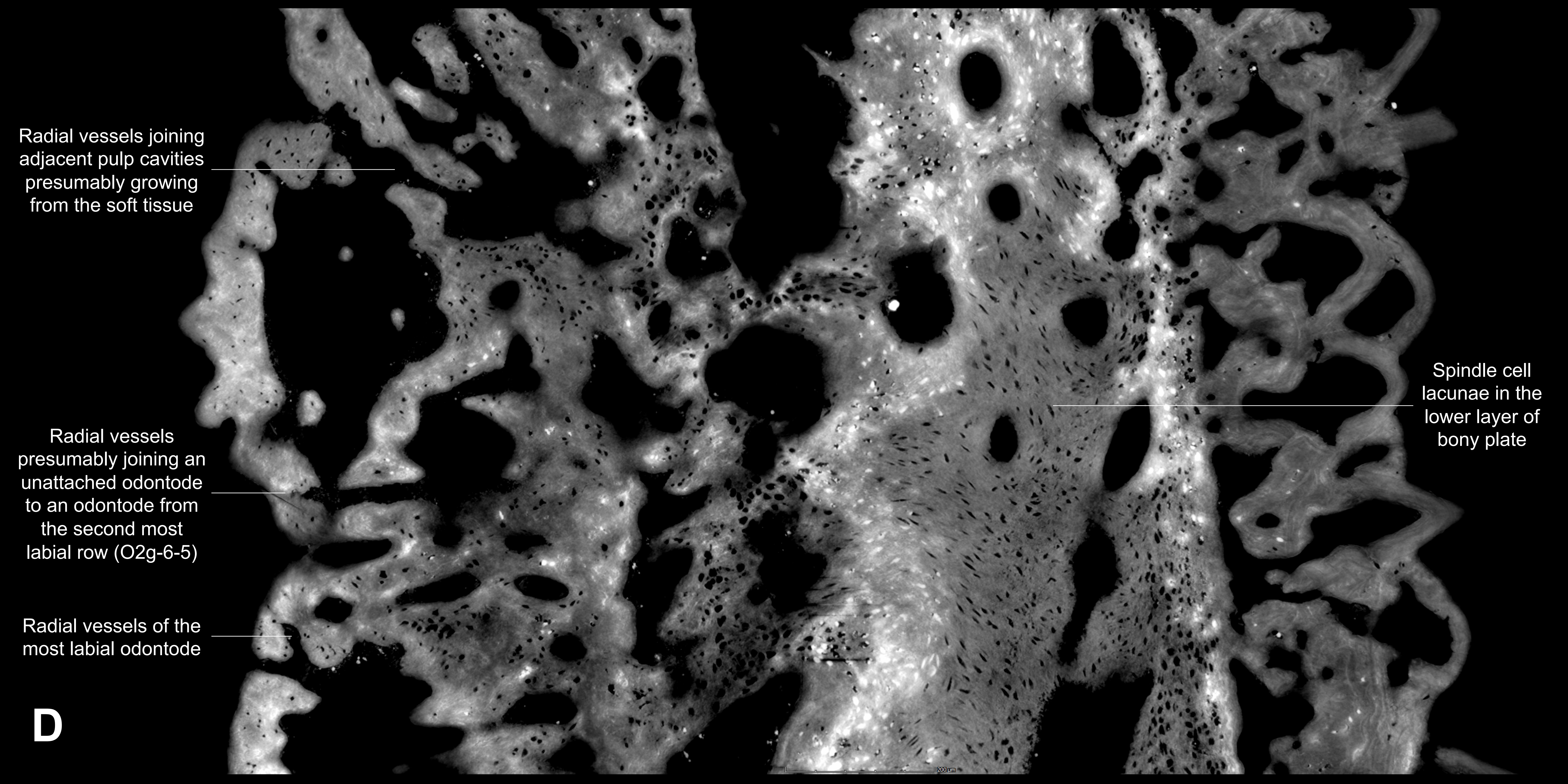

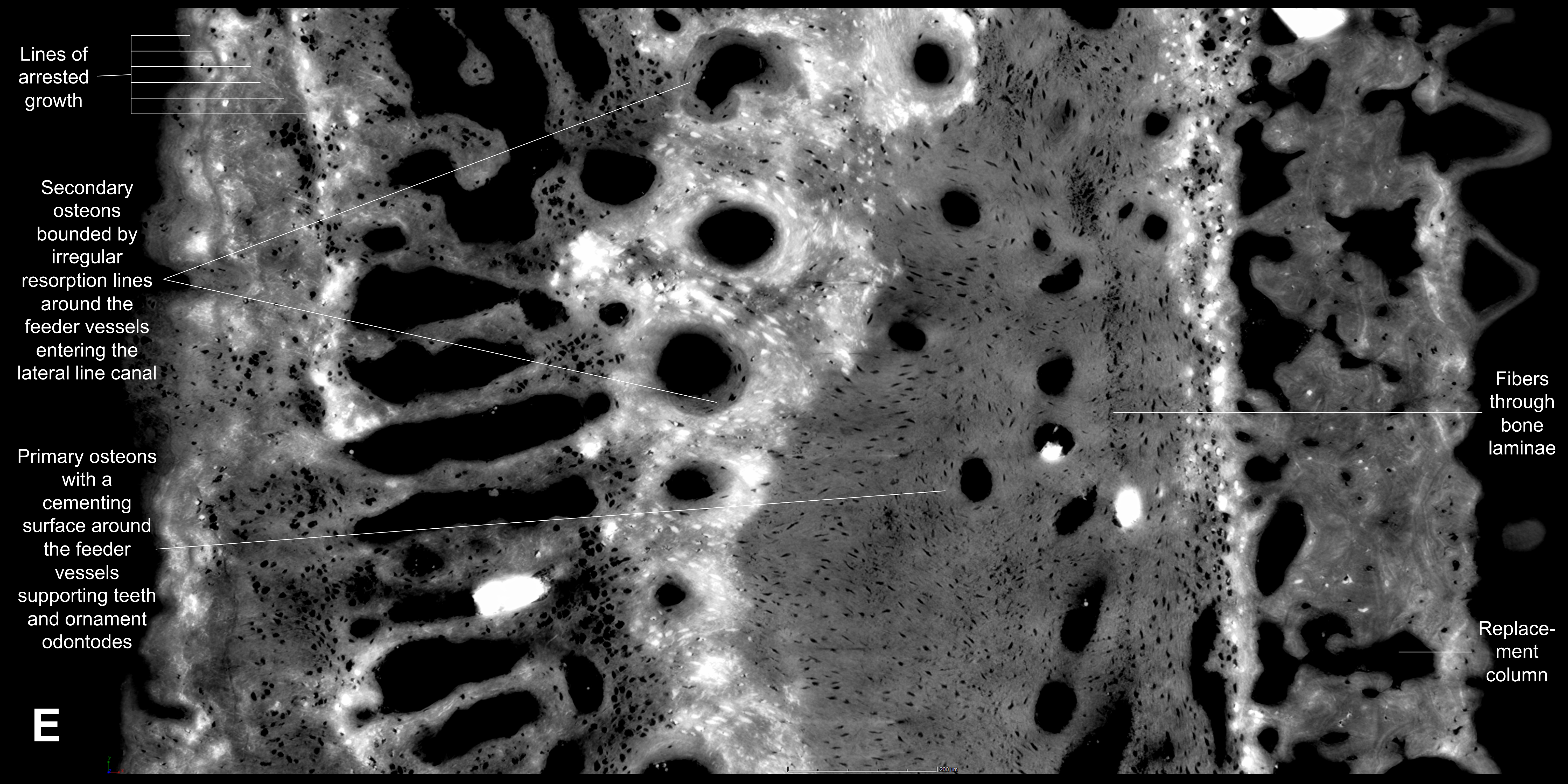
